# Extrafollicular activation of B cells contributes to the long-term survival of CD28KO mice infected with Sylvio X10/4*Trypanosoma cruzi* parasites

**DOI:** 10.1101/2025.02.14.638268

**Authors:** Rogério Silva do Nascimento, Claudio R. F. Marinho, Érika Machado de Salles, Maria Regina D’Império Lima, José M. Alvarez

**Author notes:** These authors contributed equaly.

## Abstract

In the initial phase of infections, prior to the development of the follicular antibody response, an extrafollicular B cell response develops in lymphoid organs, with activation of cells displaying B cell receptors with low affinity to the different epitopes of the invading pathogen. To evaluate if this response allows early protection in the murine host, we investigated its role during infection with Sylvio X10/4 (a low virulence *Trypanosoma cruzi* parasite) of CD28KO mice, a mouse strain that does not develop a follicular antibody response, in parallel with infected C57BL/6 mice (used as control). We observed that *T. cruzi* infection allowed CD28KO mice to survive for a long period of time, ranging from a few weeks to almost a year. However, the CD28KO mice showed higher levels of subpatent parasitemia than C57BL/6 mice, as well as more intense and long-lasting inflammatory infiltrates in the heart. Following the CD28KO mice throughout the infection, we found a significant reduction in the number of CD8^+^ T cells in the spleen, but not CD4^+^ T cells, as well as a strong fluctuation in the number of B cells. Meanwhile, although the production of IFN-γ in the spleen was not affected by CD28 deficiency, a reduction in IL-2 production was observed in anti-CD3 stimulated CD4^+^ T cells from infected CD28KO mice. Regarding the humoral response, after confirming that the T follicular helper (TFH) cells and germinal center B cells (GC-B) were absent in the infected CD28KO mice, we demonstrated that the extrafollicular antibodies, present in CD28KO infected mice, allowed an early protection in the murine host, since the transfer of serum from chronic CD28KO mice to naïve CD28KO mice, which were infected with *T. cruzi* 24 hours later, promoted a significant reduction in parasitemia. These results demonstrate that low-affinity anti-*T cruzi* extrafollicular antibodies contribute to the control of circulating parasites, allowing Sylvio X10/4-infected CD28KO mice to survive for a long time.

**AUTHOR SUMMARY:** Low-affinity anti-*T cruzi* extrafollicular antibodies contribute to the control of circulating parasites, allowing Sylvio X10/4-infected CD28KO mice to survive for a long time.

## INTRODUCTION

Chagas disease (American trypanosomiasis) described in humans by Carlos Chagas [1], is a zoonosis caused by the protozoan *Trypanosoma cruzi*, that affects around 6 million people, mostly in Latin America [2]. Human infection by *T. cruzi*, an obligate intracellular parasite transmitted by hematophagous reduviid insects [3], most often occurs after the infected insect has fed on a blood meal depositing its contaminated feces near the bite site, but can also occur through oral, transplacental, and other less common routes [2,4]. The natural course of Chagas disease includes an initial acute phase, characterized by the presence of parasites circulating in the bloodstream, and a chronic phase with subpatent levels of parasitemia. The acute phase may be asymptomatic, or present symptoms that can be confused with a cold. However, 5% of cases, mostly newborns and children, develop generalized adenopathy and hepatosplenomegaly, with myocarditis and meningoencephalitis that can lead to death [4]. Chagas disease has no spontaneous cure, the parasite persists in low numbers throughout the host’s life [5]. Remarkably, it is in the chronic phase that its most devastating consequences occur, with the development of cardiomyopathy, which affects about 30% of infected individuals, and of mega-esophagus and/or megacolon in 10-15%.

In the mammalian host, the extracellular forms of *T. cruzi* named trypomastigotes invade several types of cells, including macrophages and cardiomyocytes. In the cytoplasm, the parasite escapes the phagolysosomal vacuole and differentiates into amastigotes, which proliferate through several cycles of binary fission. Then, the amastigotes differentiate into trypomastigotes, which lyse the host cell to invade other cells [3].

Despite not achieving complete elimination of *T. cruzi*, the immune response plays an essential role in controlling the parasite. Activation of naive CD4^+^ and CD8^+^ T cells specific for *T. cruzi* requires recognition by T cell receptors of parasite peptides displayed on major histocompatibility complex (MHC) molecules on the surface of professional antigen-presenting cells (APCs), as well as interaction of CD28 molecules on T cells with B7.1/B7.2 costimulatory molecules expressed on parasite-presenting APCs. The delivery of these signals promotes the expansion of parasite-specific T cells, a process that requires the release of interleukin 2 (IL-2) and the expression of high affinity receptors for this cytokine on the surface of activated T cells [6].

One of the most important functions of activated CD4^+^ T cells is to allow the development of a humoral response by B cells, that is, the production of specific antibodies against the parasite. Among these, IgG antibodies are crucial in the control of circulating trypomastigotes, where antigen-IgG complexes act mainly by activating the complement system, a process that notably improves the phagocytic activity of macrophages [7]. In addition to allowing the development of the humoral response by B cells, CD4^+^ T cells contribute, together with CD8^+^ T cells, to the production of IFN-γ, a macrophage-activating cytokine that optimizes the destruction of phagocytosed trypomastigotes [8]. Furthermore, as amastigotes replicate intracellularly in a wide range of cells, including macrophages and cardiomyocytes, cytotoxic CD8^+^ T cells are fundamental tools for the elimination of *T. cruzi*-infected cells [9].

In acute murine infection by *T. cruzi* and other parasites, extensive activation of B cells occurs, resulting in the production of immunoglobulins with a wide range of reactivities including autoantibodies [10–11]. This broad activation of B cells may result from an intense extrafollicular response that occurs in situations such as acute parasitic infections, where a large amount of diverse (self and non-self) antigens are available. It is a CD4^+^ T cell-dependent process [12], which includes immunoglobulin class switch, although its details have not yet been fully elucidated. As evidence of being a physiological process, concomitantly with the extrafollicular B cell response, the T-B cell interaction slowly consolidates the production of high-affinity antibodies to the parasite. This process depends on the differentiation of CD4^+^ T cells into TFH cells and their interaction with B cells, which undergo a mutation process in the germinal centers followed by selection of those lymphocytes that have the greatest affinity for parasite antigens [13].

*T. cruzi* parasites are highly heterogeneous in terms of genotype, phenotype and pathogenicity [14]. In a previous study, CD28KO mice were shown to be highly susceptible to infection by the highly virulent *T. cruzi* parasites of the Y strain [15], resulting in the death of the animals a few weeks after infection. As a counterpoint, our group showed that CD28KO mice survived for an extended period of time when infected by low virulence Sylvio X10/4 *T cruzi* parasites [16], a clone of *T. cruzi* derived from a patient with acute chagasic cardiomyopathy [17]. Although infection by Sylvio X10/4 parasites does not lead to overt levels of parasitemia, it exibits marked myotropism, and in certain strains of mice it induces chronic cardiac lesions similar to those observed in humans [18]. In the present work we have analyzed in detail the behaviour of CD28KO mice infected by Sylvio X10/4 parasites, aiming to understand the physiological role played by the extrafollicular response of B cells.

## MATERIAL AND METHODS

### Sylvio X/10/4 *T. cruzi* parasites

Trypomastigotes of *T. cruzi* clone Sylvio X10/4 [19–20] were maintained by weekly passages in monolayers of LLC-MK2 cells (American Type Culture Collection – ATCC-CCL7.1). On the 7th day of culture, the day of the parasite’s ascending peak, the supernatant was collected, centrifuged at 4°C, 1200 rpm for 7 minutes, and the parasite pellet resuspended in RPMI1640 culture medium (Gibco).

### Mice and infection

Six-to-eight weeks old CD28KO and C57BL/6 (originally from The Jackson Laboratory) female mice bred under specific pathogen-free conditions at the Isogenic Mice Facility (Instituto de Ciências Biomédicas/Universidade de São Paulo, São Paulo, Brazil) were infected with Sylvio X10/4 parasites or left uninfected. For most experiments, the mice were inoculated intravenously (i.v.) with 2 x 10^5^ parasites. However, for the evaluation of the best protocol in our studies, mice were inoculated either intraperitonealy (i.p.) with 1 x 10^5^ or i.v. with 2 x 10^5^ parasites. Moreover, for the analysis of TFH cells and GC-B cells, mice were infected i.v. with 5 x 10^6^ parasites and treated with the anti-parasite drug benzonidazole (LAFEPE, Brasil), administered by gavage on day 7 of infection, in a single dose of 1g/kg of body weight, to reduce the parasite load and increase the mouse survival time.

### Subpatent parasitemia analysis

The subpatent parasitemia were evaluated from the beginning of the infection to different moments at the chronic phase. For this, the culture test in LIT (liver infusion tryptose) medium was used, a procedure where the number of epimastigotes generated from blood samples of *T. cruzi*-infected mice is estimated. Briefly, samples of 1 μL and 5 μL of blood obtained from the tail of each infected mouse were incubated in LIT medium [21]. Blood samples were deposited (in quintuplicate) in 48-wells culture plates containing 1 ml of LIT medium, incubated at 28°C and monitored after 25 days to assess the number of epimastigotes. The examination of the cultures was carried out in an inverted microscope according to a previously established standard.

### Histopathological analysis

Mouse hearts were fixed in 10% buffered formaldehyde and embedded in paraffin. Serial 3 μM sections were stained with hematoxylin and eosin for histopathological analysis. Results were expressed according to an inflammatory lesion score, where animals with greater inflammation received the score 5 and 0 the lowest one. Results were expressed as the arithmetic mean of individual scores of animals in the same group.

### Flow cytometry analysis

Spleen cells were analyzed by flow cytometry. Aliquots of cell suspensions (obtained by mechanical disruption of the spleen in RPMI medium - 3% FBS) were incubated with antibodies against different surface (extracellular labeling) and intracellular (intracellular labeling) molecules. Cells were labeled with monoclonal antibodies to mouse CD4 (H129.9), CD8 (53-6.7), CD19 (1D3), CD138 (281-2), FAS (Jo2), GL-7, PD-1 (J43), CXCR5, and IFN-γ (XMG1.2), conjugated to fluorocromes FITC (Fluorescein Isothiocianate), PE (Phicoeritrin), PE CY5 (Cy cycrome), PERCP (Peridin Clorophill Protein), Alophicocyanin (APC,), & Pacific Blue, obtained from Pharmigen (San Diego, CA). The frequency of IL-2-secreting CD4^+^ and CD8^+^ cells was estimated by capturing this cytokine immediately after its autocrine production by these cells, following the manufacturer’s protocol (BD Biosciences). Splenocytes (2×10^6^ cells/well) were placed in ice-cold media for 10 minutes with a double capture antibody that recognizes both IL-2 and the CD45 lymphocyte surface antigen. Then, cells were left unstimulated, or stimulated at a temperature of 37°C, 5% CO_2_, for 45 minutes with anti-CD3 antibody, in a continuous rotating apparatus (Macs Mix – Miltenyi Biotec). After Tstimulation, cells were labelled with fluorochrome-labelled anti-CD4 (H129.9), anti-CD8 (53-6.7) and anti-IL-2 antibodies (Pharmingen), as well as with a cell viability marker (Life Technologies). Paraformaldehyde-fixed samples were analyzed on FACS Canto or LSRF Fortessa (Becton Dickinson).

### Antibody and cytokine detection by ELISA

The presence of *T. cruzi*-specific IgM and IgG2C antibodies in sera from infected mice and uninfected controls was estimated by ELISA. Antibodies, including conjugates and standards, were purchased from Southern Biotechnologies Associates (ThermoFischer, USA). In addition, spleen cell supernatants were evaluated for cytokine determination by Capture ELISA, according to the manufacturer’s recommendations (R&D Systems, USA) and compared to standard recombinant cytokines curves.

### Statistical Analysis

For comparative analysis between the experimental groups, the Student t test for two variables assuming equal variance and the ANOVA test for multiple comparison were used. Significance levels were defined as being less than or equal to 0.05 (p<0.05).

## RESULTS

### Survival curve of CD28KO mice along infection with *T. cruzi*

To demonstrate the impact of CD28 deficiency on Sylvio X10/4 parasite infection, Figure 1A illustrates the survival curves of CD28KO and C57BL/6 mice following i.p. infection with 1 x 10^5^ parasites or i.v. infection via the ophthalmic plexus with 2 x 10^5^ parasites. The graphs show that, regardless of the route or inoculum size, CD28KO mice had greater susceptibility to Sylvio X10/4 infection compared to C57BL/6 mice. Notably, CD28KO mice succumbed to infection at varying intervals over a prolonged period, ranging from day 40 to day 245 post-infection (p.i.). Additionally, in CD28KO mice, the i.v. infection with 2 x 10^5^ parasites confered prolonged survival compared to i.p. infection with 1 x 10^5^ parasites and was used in subsequent experiments.

**Figure 1.**
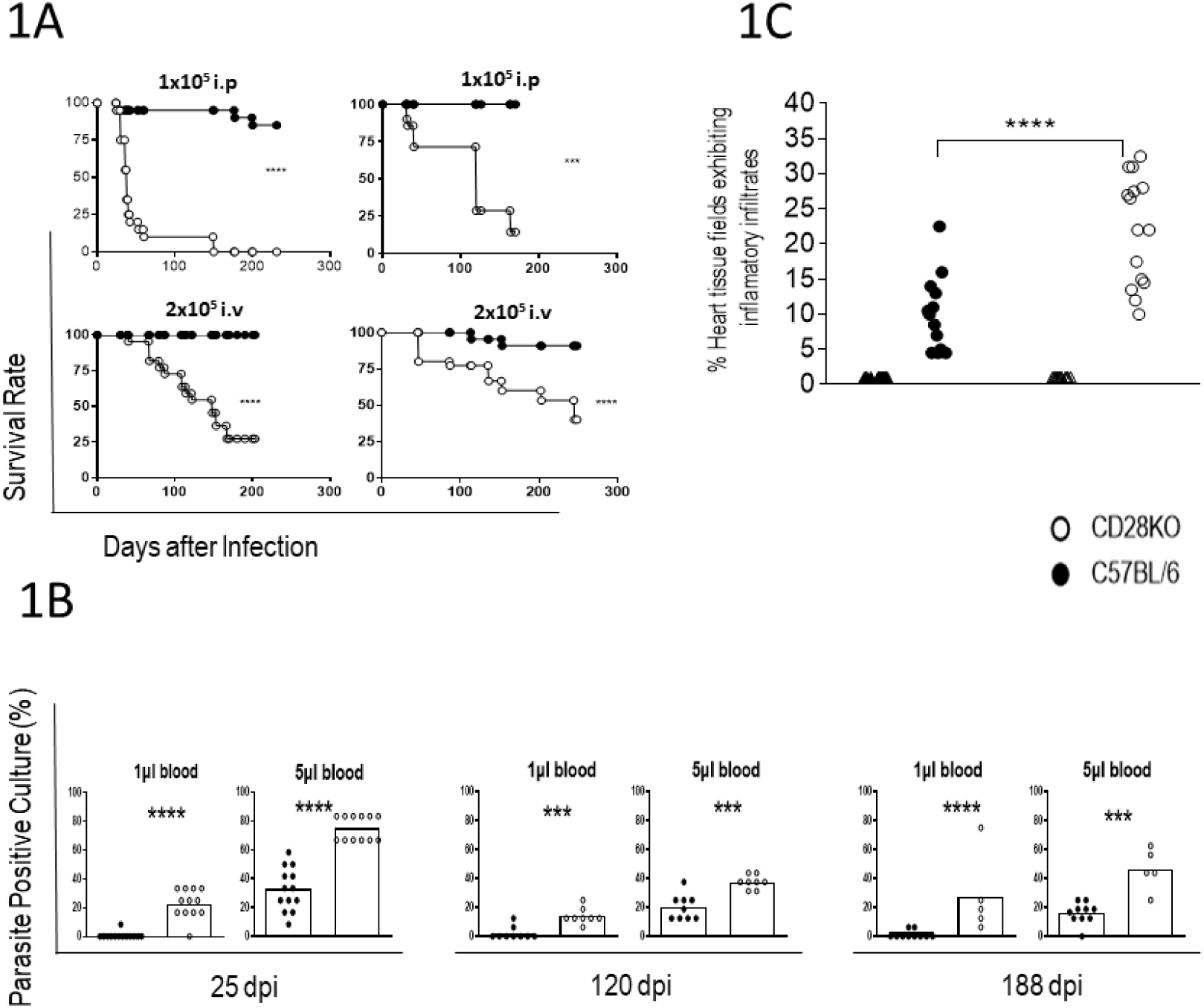
Survival curve, parasitemia and heart pathology of CD28KO and C57BL6 mice infected with Sylvio X10/4 parasites. **A)** Two representative experiments showing the survival curves of CD28KO and C57BL/6 mice infected i.p. with 1 x 10^5^ Sylvio X10/4 parasites (above), or i.v. with 2 x 10^5^ Sylvio X10/4 parasites (below). Student’s-t-test. ***, P<0.001; ****, P < 0.0001, for plots with groups of infected CD28KO and C57BL/6 mice. **B)** Relative subpapent levels of parasitemia in CD28KO and C57BL/6 mice along i.v. infection with 2 x 10^5^ Sylvio X10/4 parasites. Blood aliquots of 1 and 5 microliters obtained from each mouse in the same experiment, at the acute-chronic interphase (27 days p.i.), and at two later moments of infection (120 days p.i. and 188 days p.i.), were cultured in quintuplicates in LIT medium and screened by microscopy for epimastigote growth. Frequency of positive cultures are shown. Student’s-t-test. **, P<0.01; ***, P<0.001; ****, P < 0.0001 between groups of infected C57BL/6 and CD28KO mice on the same plot. **C)** In compilated experiments, C57BL/6 and CD28KO mice chronically-infected with 2 x 10^5^ Sylvio X10/4 parasites, and respective uninfected controls, were sacrificed and cardiac tissue slides stained with H/E (hematoxylin/eosin) for microscopic evaluation. Each point in the graphs refers to an arbitrary inflammatory score of individual mice, obtained by estimating the frequency of fields exhibiting inflammatory infiltrates in the different regions of the cardiac tissue. Statistics for compiled experiments are shown. One Way Anova Test. Groups: C57BL/6 control (black triangle), CD28KO control (empty triangle), infected C57BL/6 (black circle), infected CD28KO (empty circle).

### Subpatent parasitemia levels of Sylvio X10/4 parasites in CD28KO mice compared to C57BL/6 mice

To assess the relative levels of subpatent parasitemia in CD28KO and C57BL/6 mice along the i.v. infection with 2 x 10^5^ Sylvio X10/4 parasites, 1 and 5 µL blood samples obtained at different time points from infected mice were cultured in LIT (liver infusion tryptose) medium for 25 days. After the incubation period, the number of epimastigotes was quantified as an indirect measurement of parasitemia. These are the extracellular proliferative forms of *T. cruzi*, present in the gut of the insect vector, which are generated *in vitro* when non-dividing blood trypomastigotes are incubated in axenic cultures with LIT medium. Using this approach, we investigated whether chronically infected mice displayed varying levels of subpatent parasitemia over the course of infection. For this, aliquots of blood obtained from CD28KO and C57BL/6 mice on days 27 (acute/chronic interphase), 120 and 188 (late chronic phase) of infection with 2 x 10^5^ Sylvio X10/4 parasites were cultured in LIT medium and screened by microscopy for epimastigote growth. Samples derived from infected CD28KO mice always exhibited higher levels of subpatent parasitemia than those from infected C57BL6 mice.

Interestingly, in the infected CD28KO mice slightly higher parasitemias were observed at day 27 compared to those at days 120 and 188, while no apparent differences were observed between the two chronic time points (Figure 1B).

### Histopathological analysis of the heart of CD28KO mice infected with Sylvio X10/4 parasites

Because heart pathology is a major impact of *T. cruzi* infection, we evaluated the intensity of inflammatory infiltrates in this organ (atrium and ventricle) in CD28KO and C57BL/6 mice along the course of i.v. infection with 2 x 10^5^ Sylvio X10/4 parasites. The objective was achieved by counting the extent of infiltrates in heart slides of individual mice.

CD28KO mice developed more extensive inflammatory infiltrates in the heart compared to those observed in infected C57BL/6 mice (Figure 1C, that shows a compendium of data from mice sacrificed on days 64 to 359). Nests of *T. cruzi* amastigotes were not found by microscopy examination in heart slides of both infected groups along the course of the infection.

Based on the intensity of the cardiac infiltrates, our results indirectly indicate that, compared to the C57BL/6 mice, CD28KO mice suffer a greater invasion of cardiac tissue by *T cruzi* Sylvio X10/4 parasites and/or display a lower capacity to eliminate them from this organ.

### Spleen cellularity in *T. cruzi*-infected CD28KO and C57BL/6 mice

The spleen is a secondary lymphoid organ which responds to microorganisms circulating in the blood with activation and proliferation of specific B and T lymphocytes. Compared to non-infected mice, both C57BL/6 and CD28KO mice infected with 2 x 10^5^ Sylvio X10/4 parasites showed a considerable increase in the total number of spleen lymphocytes in the acute phase (day 13 p.i.), that declined in the chronic phase, when it became closer to that in mice from the non-infected groups (Figure 2A, left). Interestingly, at day 13 p.i., despite the increase in the spleen lymphocyte numbers, there was a relative reduction in the frequency of these cells in both mouse strains (Figure 2A, right).

**Figure 2.**
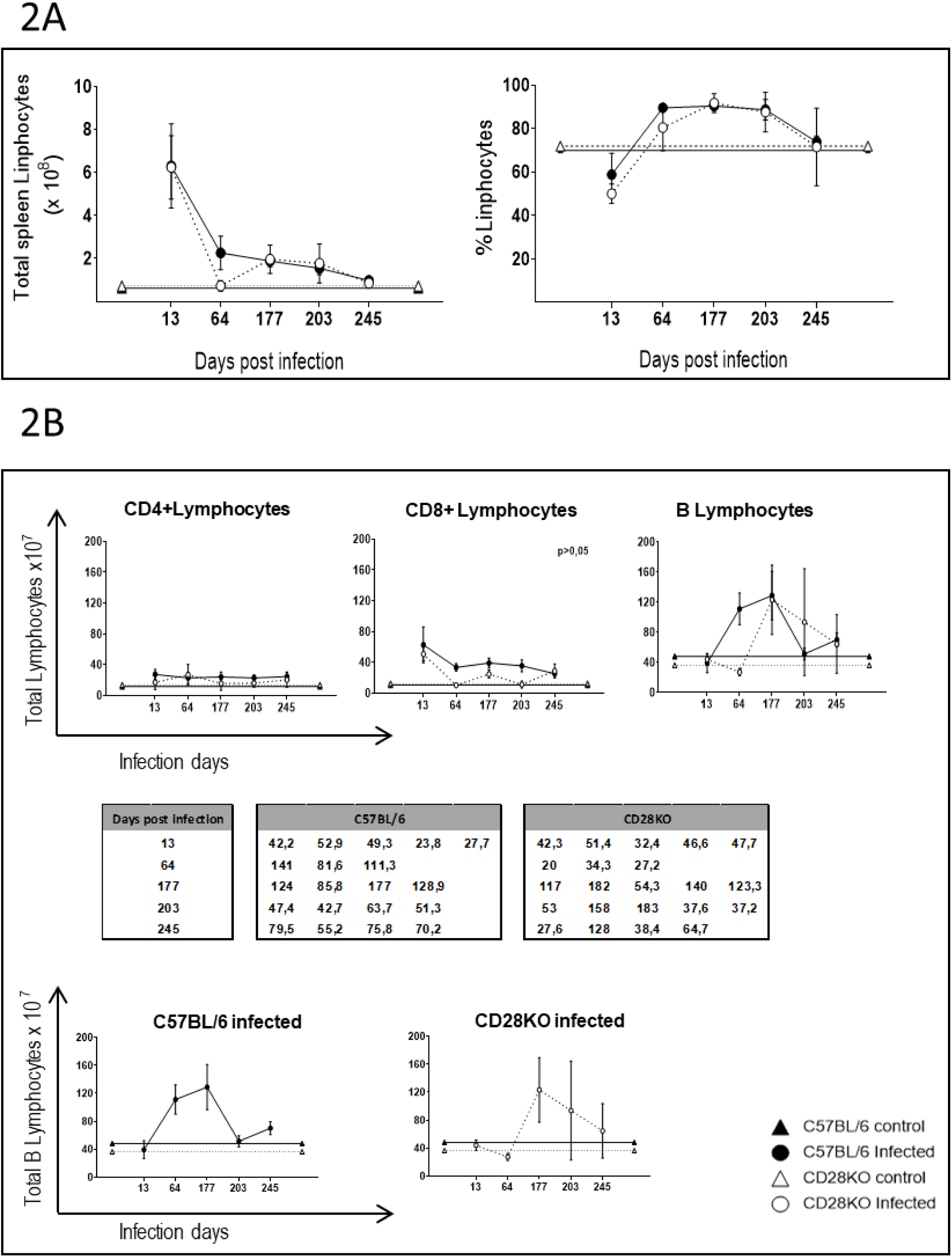
Frequency and total number of CD4^+^ and CD8^+^ T lymphocytes, and B (CD19^+^) lymphocytes, in the spleen of CD28KO and C57BL/6 mice infected with Sylvio X10/4 parasites. Data represent a single experiment, carried out from days 13 to day 245 p.i. in groups of i.v. infected CD28KO and C57BL/6 mice and corresponding non-infected groups. The mice were sacrificed for spleen processing and analysis by flow cytometry, with the total number of cells in the spleen of individual mice calculated as indicated in the Material and Methods. **A)** Total number (left) and frequency (right) of lymphocytes. **B)** ABOVE: Total number of CD4^+^ and CD8^+^ T lymphocytes, and B (CD19^+^) lymphocytes. MIDDLE and BELOW: The variability of the total number of B lymphocytes from individual mice at different time points of infection is also shown separately (both as a table and as graphs), for CD28KO and C57BL/6 mice, in order to highlight the heterogeneity of B cell numbers present in CD28KO mice. Test *ANOVA*, p<0.05 for the plots of CD8^+^ T cells between the groups of C57BL/6 and CD28KO infected mice.

### Analysis of T and B cell populations in the spleen of Sylvio X10/4-infected CD28KO and C57BL/6 mice

Over the course of infection, when the total number of lymphocytes gradually decreases in the spleen of infected mice, a small, non-significant, lower number of CD4^+^ T cells was found in CD28KO mice compared with C57BL/6 mice. Meanwhile, a significant reduction in the total number of CD8^+^ T cells was observed in CD28KO mice over the course of infection, as well as a broad individual heterogeneity in the number of B lymphocytes (Figure 2B, above). To more clearly visualize this last issue, the total number of B lymphocytes at the different time points of infection is also shown separately for the two mouse strains, a procedure that highlights the individual variation in the number of B cells present in chronic CD28KO mice (Figure 2B, middle and below). ANOVA test: ns, not significant (middle chart and bottom figures).

### Analysis of IL-2 production by spleen cells of CD28KO and C57BL/6 mice infected with Sylvio X10/4 parasites

To verify the impact of CD28 deficiency on IL-2 production, a parameter related to the proliferation of T cells, we analyzed the frequency of IL-2-producing CD4^+^ and CD8^+^ T lymphocytes from CD28KO and C57BL/6 mice infected i.v. for 15 days with 2 x 10^5^ Sylvio X10/4 parasites, as well as in those from non-infected controls (Figure 3A). The frequency of IL-2-secreting CD4^+^ or CD8^+^ T cells was evaluated by capturing IL-2 immediately after its autocrine production, as described in the Material and Methods. In the absence of *in vitro* re-stimulation with anti-CD3 antibodies, a significantly higher frequency of IL-2-producing cells was observed among CD8^+^ T lymphocytes from infected CD28KO mice (Figure 3A, right), whereas it did not reach significance among CD4^+^ T lymphocytes (Figure 3A, left). These results suggest that in the absence of CD28, the long-term production of IL-2 in response to *T. cruzi* is largely due to CD8^+^ T cells. Moreover, the higher basal frequency of IL-2-producing CD4^+^ and CD8^+^ T cells in CD28KO mice infected for 15 days (compared to infected C57BL/6 mice) suggests that, at this time point, T cell replication is still ongoing in infected CD28KO mice but has already declined in infected C57BL/6 mice.

**Figure 3.**
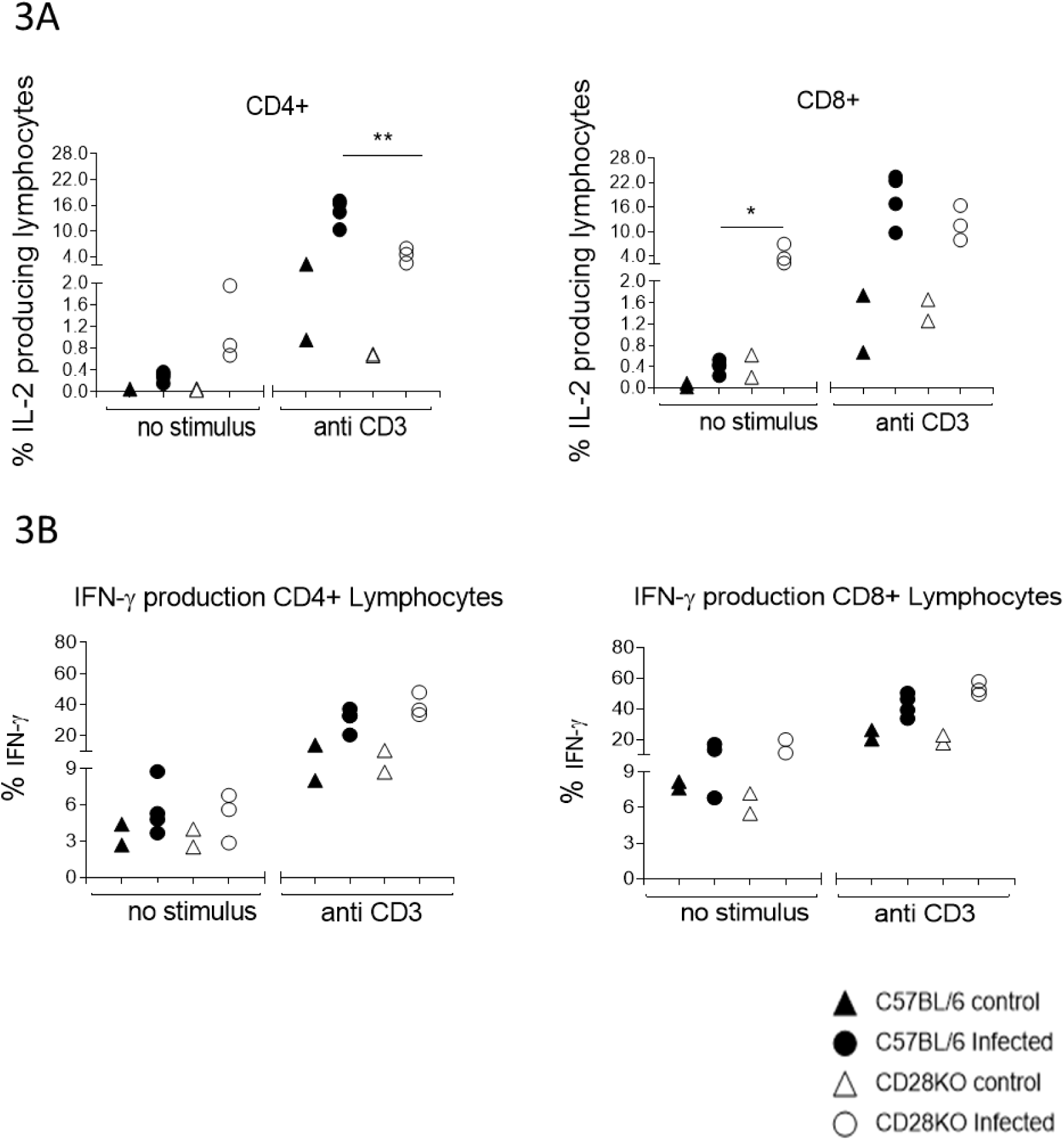
IL-2 and IFN**γ** production by CD4^+^ and CD8^+^ T lymphocytes on day 15 p.i. (acute phase) by Sylvio X10/4 parasites, in the absence of re-stimulation and after *in vitro* stimulation with anti-CD3 antibodies. A) **Representative dot-blots showing** the frequency of IL-2 producing CD4^+^ T cells (left Figure), and CD8^+^ T cells (right Figure) in the spleen of CD28KO and C57BL/6 mice infected for 15 days with 2 x 10^5^ Sylvio X10/4 parasites and corresponding uninfected controls, in the absence of *in vitro* stimulation, and after a secondary *in vitro* overnight stimulation with anti-CD3 antibodies for 6 hours. Non-parametric T-test comparing the infected CD28KO and C57BL/6 groups. *, P<0.05; **, P<0.01; and ns, not significant**. B)** Representative graphs of IFNγ-producing CD4^+^ and CD8^+^ T spleen lymphocytes from uninfected and Sylvio X10/4-infected CD28KO and C57BL/6 mice that were cultured for six hours, without stimulation (left side graphs) or after *in vitro* stimulation with anti-CD3 antibodies (right side graphs). C57BL/6 control mice (black triangle), infected C57BL/6 mice (black circle), control CD28KO mice (empty triangle) and infected CD28KO mice (empty circle). The graphs indicate the percentage of IFNγ−producing cells in the spleen of C57BL/6 and CD28KO mice on day 15 of infection with 2 x 10^5^ Sylvio X10/4 parasites and respective uninfected controls. Mean (control C57BL/6 mice, n = 2; infected C57BL/6 mice, n = 4; control CD28KO mice, n = 2; infected CD28KO mice, n = 3). Nonparametric T test comparing infected CD28KO mice with infected C57BL/6 mice group. ns, not significant.

In contrast, after *in vitro* stimulation with anti-CD3 antibodies, significantly lower IL-2 production was observed among CD4^+^ T cells of CD28KO mice, while no difference was observed among CD8^+^ T cells compared to that of infected C57BL/6 mice. These results support the hypothesis that in acutely-infected mice, the absence of CD28 primarily impacts the IL-2 response of CD4^+^ T lymphocytes.

### Analysis of IFN-**γ** production by CD4^+^ T and CD8^+^ T spleen cells of CD28KO and C57BL/6 mice infected with Sylvio X10/4 *T cruzi* parasites

IFNγ producing CD4^+^ T (Figure 3B left) and CD8^+^ T (Figure 3B right) spleen lymphocytes from CD28KO and C57BL/6 mice were analyzed on the 15 day p.i. Our results show that, in both lymphocyte populations, equivalent levels of this cytokine were produced by infected CD28KO and C57BL/6 mice. Furthermore, similar results were also observed in both groups after *in vitro* stimulation with anti-CD3 antibodies. However, in all experiments, the in *vitro* production of IFNγ by non-stimulated CD4^+^ and T CD8^+^ T cells from *T. cruzi*-infected and control mice were considerably lower than those of corresponding anti-CD3-stimulated cells.

### Infection of CD28KO mice by Sylvio X10/4 parasites does not induce the formation of splenic TFH cells

To confirm that in *T. cruzi*-infected CD28KO mice no other source of antibodies occurred besides the extracellular ones, we investigated for the presence of TFH cells and GC-B cells in CD28KO mice. To analyze the presence of TFH cells by flow cytometry, spleen cells from CD28KO and C57BL/6 mice infected with 5 x 10^6^ Sylvio X10/4 parasites were stained for CD4, PD-1 and CXCR5 to identify the TFH cell population (CD4^+^PD-1^High^CXCR5^High^). As indicated above, and in the Material and Methods section, a high parasite dose was necessary to clearly visualize this subset by flow cytometry. Our experiments showed that, while in infected C57BL/6 mice the TFH population was clearly visible, being measurable from day 14 to 25 p.i. (data not shown), in infected CD28KO mice there was no generation of these cells (Supplementary Figure 1A)

### Infection of CD28KO mice with Sylvio X10/4 does not induce splenic GC-B cells

Moreover, to evaluate by flow cytometry the presence of GC-B cells in the spleen of C57BL6 and CD28KO mice infected by Sylvio X10/4 parasites, splenocytes were stained for CD19 (for B lymphocytes analysis), CD138 (for plasmocytes analysis), FAS and GL-7, to characterize the subpopulation of GC-B lymphocytes that have the CD19^+^CD138^-^GL-7^High^Fas^High^ phenotype. As indicated for the evaluation of TFH cells, in the experiments with GC-B cells, the mice were infected with a high dose of Sylvio X10/4 parasites for a clear generation/visualization of GC-B cells in the C57BL/6 mouse. We found that CD28KO mice infected i.v. with 5 x 10^6^ Sylvio X10/4 parasites do not generate GC-B cells, while those of infected C57BL/6 mice generate an expressive subpopulation of these lymphocytes (Supplementary Figure 1B).

Our results indicate that, unlike the C57BL/6 mice, mice deficient in the costimulatory molecule CD28 do not have the capacity to generate TFH cells and GC-B cells following infection with Sylvio X10/4 parasites.

### Infection of CD28KO mice by Sylvio X10/4 parasites induces production of IgM and IgG antibodies against the parasite

The humoral immune response includes the activation of both extrafollicular and follicular B cells in humans and mice. To evaluate the humoral response to Sylvio X10/4 parasites in the absence of the costimulatory molecule CD28, serum samples from infected CD28KO mice were collected at the acute phase (day 16 p.i.), and during the chronic phase (days 47 and 69 p.i.). These samples, as well as those from uninfected CD28KO mice (which were used as controls), were then evaluated by ELISA for the production of anti-*T. cruzi* antibodies of the IgM and IgG classes. Sylvio X10/4 antigen was used for plaque sensitization. During the acute phase, as well as throughout the chronic phase (47-69 days p.i.), sera from infected CD28KO mice exhibited a significant IgM reactivity towards Sylvio X10/4 parasites (Figure 4A, left). Additionally, a notable higher IgG antibody response to the parasite was also observed in sera from infected CD28KO mice. However, unlike IgM, the anti-*T. cruzi* IgG antibody level was less pronounced during the chronic phase compared to the acute phase (Figure 4A, right), a result that could be due to IgG consumption both in the blood and the infected tissues.

**Figure 4:**
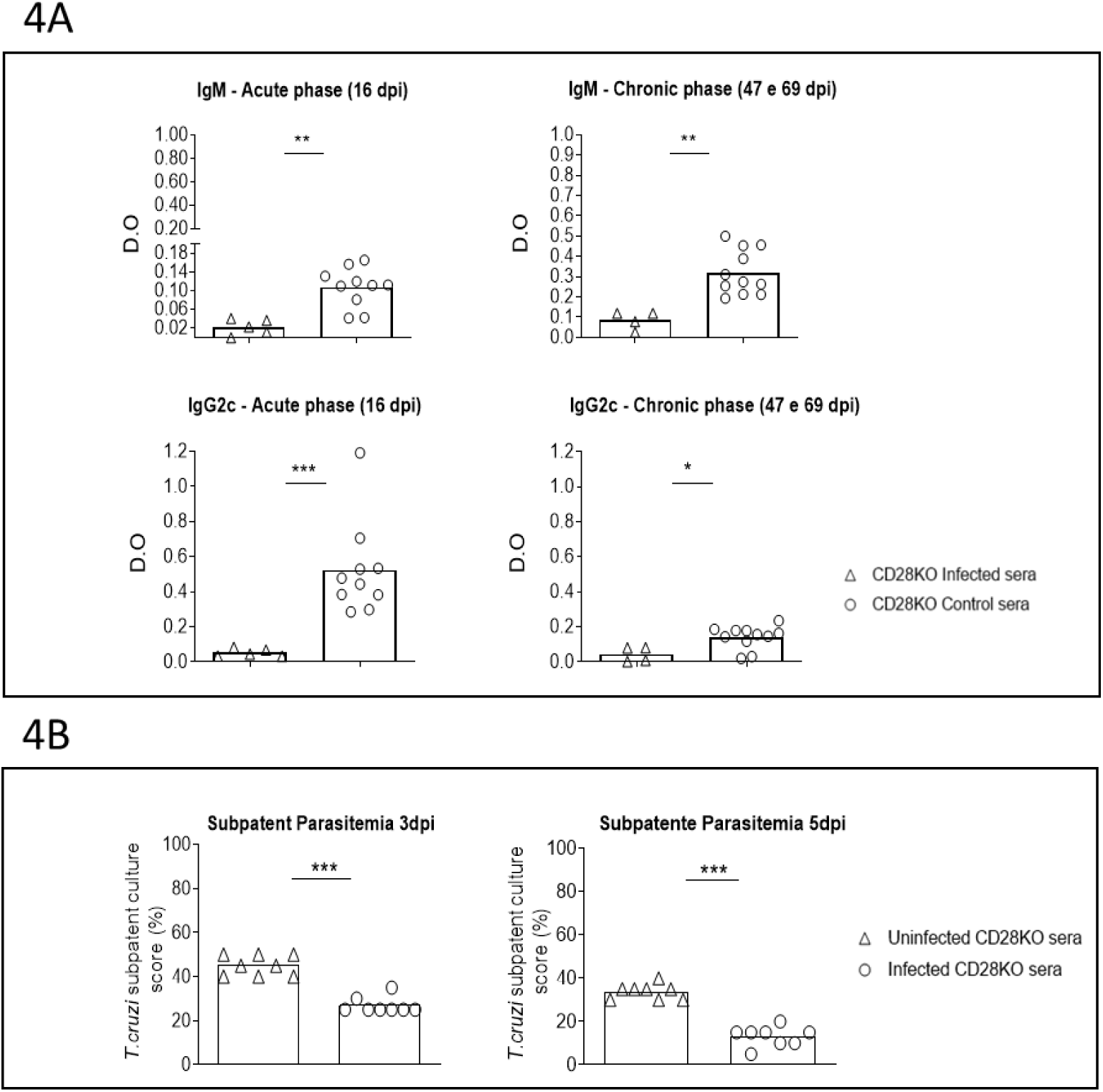
Antibody response and protection capacity in CD28KO mice to Sylvio X10/4 parasites. **A)** Serum levels of IgM and IgG antibodies against Sylvio X10/4 parasite antigen, in the acute (16 day p.i.) and chronic (47 and 69 days p.i.) phases of infection by Sylvio X10/4 parasites, compared with those in the sera from uninfected CD28KO mice (CD8KO control). The results presented in the graphs were obtained by indirect ELISA, subtracting the Optical Density (O.D.) value of each serum against the blockade, from its respective O.D. value against the Sylvio X10/4 antigen. Each point represents the arithmetic mean of triplicates. Non-parametric T test comparing the CD28KO group infected with Sylvio X10/4 with the uninfected CD28KO group. *, P<0.05; **, P<0.01; and ***, P<0.001. **B)** Naïve CD28KO mice treated i.v. with serum from CD28KO mice infected with Sylvio X10/4 parasites for 74 days, or with serum from uninfected CD28KO mice, were challenged i.v. 24hrs later with 2×10^5^ Sylvio X10/4 parasites, and their blood parasitism evaluated in the LIT assay after 3 and 5 days.

To evaluate if protection could be attained by chronic serum transfer, 100 μL of chronic serum (obtained from CD28KO mice infected for 74 days with Sylvio X10/4 parasites), or with the same volume of serum from uninfected CD28KO mice, were inoculated i.v. into naïve CD28KO mice, which were challenged i.v. 24 hours later with 2 x 10^5^ Sylvio X10/4 parasites, and screened after further 3 or 5 days for parasite level by the LIT assay. The results of these experiments showed that transfer of chronic serum from CD28KO mice conferred partial protection to naïve CD28KO mice, that could be observed when compared to that of infected mice transferred with serum from noninfected CD28KO mice (Figure 4B, left and right).

## DISCUSSION

The analysis of the B cell response in CD28KO mice infected with low virulence Sylvio X10/4 parasites is justified by the fact that these mice do not rely on the follicular antibody response to control the parasite. In consequence, this model allowed us to explore the importance of the extrafollicular B cell response, the primordial type of antibody production, that acts at the early immune response to microorganisms, and throughout life whenever the follicular antibody response is absent.

Exploring the infection of CD28KO mice with Sylvio X10/4 parasites, we observed greater survival when mice were infected i.v. with 2 x 10^5^ Sylvio X10/4 parasites than when infected i.p. with 1 x 10^5^ parasites. These results can be explained by the fact that after reaching the blood, most parasites injected i.v. go straight to the spleen, where they soon activate the extrafollicular B cell response; conversely, in i.p. inoculation, the parasite is likely to invade other cells, such as peritoneal macrophages or other tissue cells, before the adaptive immune response is initiated. Furthermore, the intensity of the extrafollicular response depends on the number of infecting parasites, hence it was stronger in the group infected with 2 x 10^5^ parasites. In view of these results, the i.v. injection protocol with 2 x 10^5^ parasites was chosen for our studies. In this regard, it is interesting to note that in CD28KO mice infected with LCMV (lymphocytic choriomeningitis virus), a virus that replicates extensively, the infecting dose does not appear to play an important role for mouse survival, whereas in VSV (vesicular stomatitis virus) infection, which replicates poorly, the viral dose becomes important [22–23], a set of results that can be interpreted as resulting from higher and lower levels of extrafollicular B cell response, respectively induced by these viruses.

Our kinetic study in mice infected i.v. with 2 x 10^5^ Sylvio X10/4 trypomastigotes, where the subpatent level of live parasites was assessed by culturing aliquots of blood in LIT medium, demonstrates that CD28 signaling is important for low virulence *T. cruzi* control, as previously described for those of high virulence [15]. Nevertheless, in our study, subpatent parasitemia levels in CD28KO mice decreased from day 27 p.i. to days 120 and 188 p.i., suggesting that some level of low virulence parasite control occurs in the absence of CD28 and permits mouse survival during a long period of time.

To understand the cause of this “prolonged survival”, we attempted to identify aspects of the immune response that are relatively preserved along the course of infection. To this end, we analyzed the kinetics of splenic lymphocytes in CD28KO and C57BL/6 mice infected with Sylvio X10/4 parasites. In these experiments, we observed that both in the acute and chronic infection the total number of CD8^+^ T lymphocytes was lower in CD28KO mice than in C57BL/6 mice. Different possibilities could explain this finding, one of which being a partial dependence on CD28 co-stimulation for the activation of these cells. Supporting this, at an early time of infection (day 15 p.i), the frequency of IL-2-producing CD8^+^ T cells in CD28KO mice was significantly higher than that observed in C57BL/6 mice, a result suggesting that their proliferation is delayed in the absence of CD28. However, as Sylvio X10/4 is a myotropic parasite, its persistence in the spleen would not be expected, indicating that the reduced presence of CD8^+^ T cells at the spleen of CD28KO mice could be mainly due to their increased migration to muscle tissues where cytoplasmic invasion by this parasite occurs. In any case, the production of IL-2 by CD8^+^ T spleen cells from infected CD28KO mice indicates that these cells are not totally dependent on CD28 signaling for IL-2 production.

On the other hand, in relation to the effector immune response at the spleen, we observed that, for both CD4^+^ and CD8^+^ T cells, the production of IFNγ was not different in Sylvio X10/4-infected CD28KO and C57BL/6 mice, an effect already described in other experimental models [24]. Therefore, among other effects, IFNγ-dependent activation of macrophages must occur in both mouse strains, ensuring the destruction by phagocytosis of IgG-opsonized trypomastigotes.

In the absence of CD28, mice infected with Sylvio X10/4 parasites did not generate TFH and GC-B cells, their B cell response being restricted to generation of extrafollicular IgM and IgG antibodies. As mentioned above, IgM suffers the limitation of being restricted to the intravascular space, the IgG antibodies being responsible for limiting tissue parasitism, in such a way that its consumption causes infected mice to lose control of the parasite. To avoid this, CD28KO mice appear to require reactivation of the extrafollicular B cell response whenever parasitism increases. This may occur in successive waves, where the level of blood parasitism dictates the intensity of the extrafollicular B cell activation, which in turn maintains blood parasitism at a safe level, allowing the infected mice to survive as long as this oscillatory balance is preserved. In this regard, in our experiments, it was interesting to observe that at the advanced chronic phase, the number of spleen B cells of individual CD28KO mice exhibited impressive variability. However, this process is occasionally insufficient to control the parasite, leading to the death of infected CD28KO mice.

Differently, whenever Sylvio X10/4 parasites invade the cytoplasm of muscle cells or other permissive cells, the invading parasites will be out of reach of B cells for 4-5 days, and, in consequence, the extracellular B cell activation will be reduced, this process changing direction when the expanded intracellular amastigotes differentiate into trypomastigotes and reach the extracellular space.

Interestingly, as expected from a persistent extrafollicular antibody response, in addition to Sylvio X10/4 parasites recognition, low-affinity reactivity against extracts from other parasites is observed in the sera of Sylvio X10/4 infected CD28KO mice (data not shown). The reactivity of sera against antigenic extracts other than the inducing parasite is probably the result of cross-reactivity, that is, the low-affinity recognition of parasites that present molecules that have epitopes similar to those of the Sylvio X10/4 parasites, a process that has been called polyclonal activation of B cells [10 and 11]. As a corollary of the studies in this manuscript, we demonstrate that the transfer of sera obtained from *T. cruzi*-infected CD28KO chronic mice to CD28KO naïve mice, confers partial protection against infection with Sylvio X10/4 parasites, a result that demonstrate the protective effect of the extrafollicular B cell response.

## Conclusion

The results of this manuscript indicate that the long-term survival of mice deficient in the costimulatory molecule CD28, infected with low-virulence Sylvio X10/4 parasites, occurs because infection with this parasite results in low (subpatent) levels of parasitemia. This situation allows partial control of the parasite by the immune system, where there is an important contribution of the extrafollicular B cell response. Importantly, we can extrapolate from the above results that the extrafollicular B cell response also operates in euthymic mice, as well as in humans, reducing parasite load at the early phase of infection.

## ACKNOWLEDGEMENTS

To FAPESP, number 2018/25984-7 and 2015/20432-8, and to CNPq, numbers 304540/2020-0, 308875/2023-0 and 3038-10/2018-1 for project funding

---

**Supplementary Figure 1.**
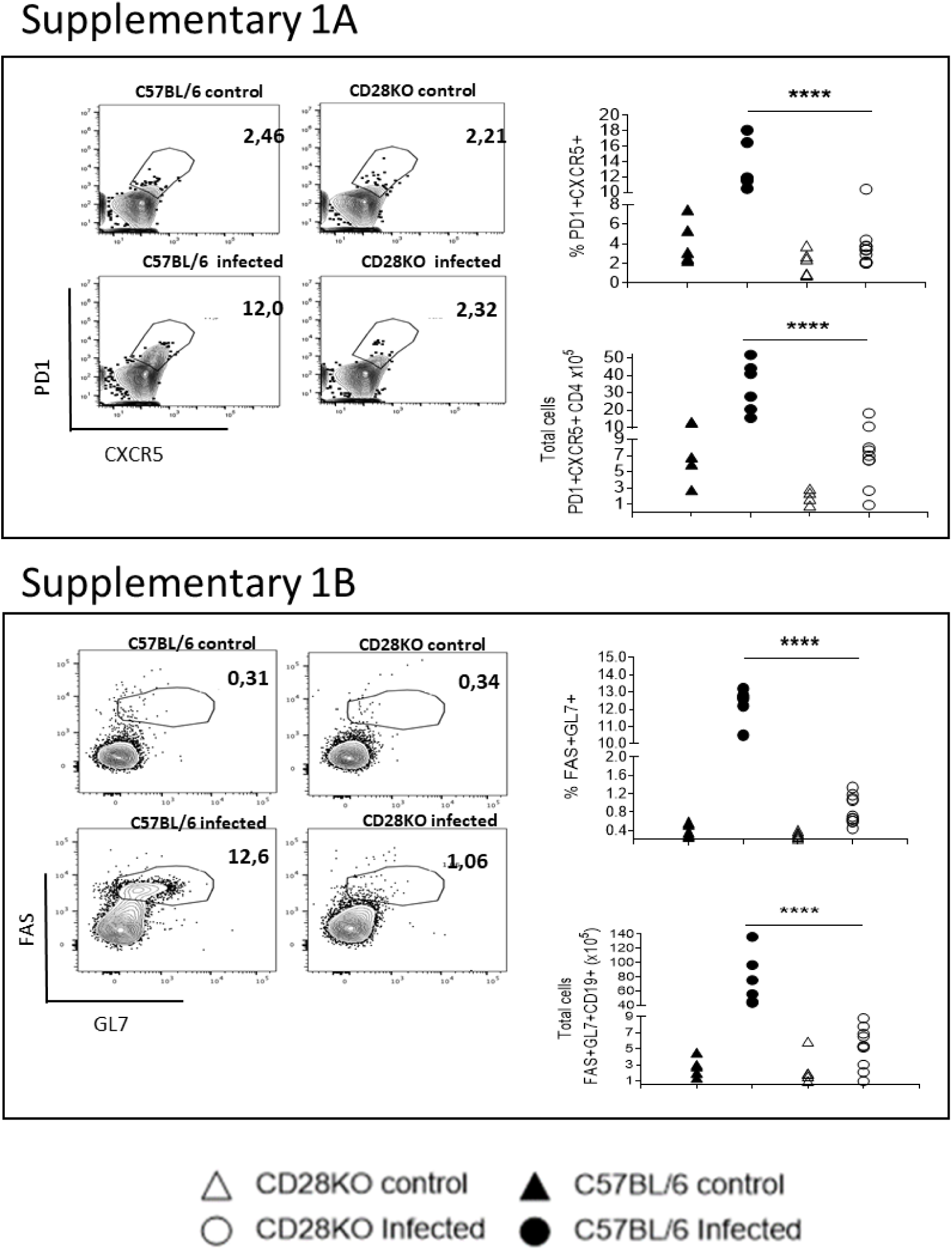
Analysis of TFH and GC-B cells in CD28KO mice infected for 16 days with Sylvio X10/4 parasites. **A)** Representative plots of TFH (CD4^+^PD-1^High^CXCR5^High^) cells in CD28KO and C57BL/6 mice infected for 16 days with 5 x 10^6^ Sylvio X10/4 parasites, and uninfected controls. The graphs show the TFH subpopulation in the infected C57BL/6 mice (lower left), while the other groups, including the infected CD28KO mice, did not clearly show this subpopulation. The graphs at the right side show the frequency and number of TFH cells in the spleen of CD28KO and C57BL/6 mice at day 16 p.i. and respective uninfected controls. Uninfected C57BL/6 mice, n = 5; infected C57BL/6 mice, n = 6; uninfected CD28KO mice, n = 5; infected CD28KO mice, n = 10). One Way Anova Test. ****, P<0.001; ns, not significant. **B)** The presence of GC-B cells, identified by the expression of FAS and GL7 markers in the B cell population (CD19^+^CD138^-^), was analyzed in the spleen of CD28KO and C57BL/6 mice on day 16 of infection with 5 x 10^6^ Sylvio X10/4 parasites, and respective uninfected controls. Dot blot figures show the presence of a GC-B cell population in infected C57BL/6 mice (lower left), which is not patent in the infected CD28KO mice (lower right) and non-infected control groups (upper figures). Graphs at the right side indicate, respectively, the frequency and number of Fas^+^GL7^+^ (GC-B cells) in CD28KO and C57BL/6 mice infected with 5 x 10^6^ Sylvio X10/4 parasites and respective uninfected CD28KO and C57BL/6 controls. The statistics of a representative experiment are shown. Mean of uninfected C57BL/6 mice, (n=5); infected C57BL/6 mice, (n=6); uninfected CD28KO mice, (n=5); and infected CD28KO mice, (n=10). Representative dot plots for GC-B cell frequency (CD19^+^CD138^-^GL7^High^Fas^High^) among spleen cells and statistics of frequencies and total numbers. One Way Anova Test. ****, P<0.001; ns, not significant.

